# Thermal sensors improve wrist-worn position tracking

**DOI:** 10.1101/552174

**Authors:** Jake J. Son, Jon C. Clucas, Curt White, Anirudh Krishnakumar, Joshua T. Vogelstein, Michael P. Milham, Arno Klein

## Abstract

Wearable devices provide a means of tracking hand position in relation to the head, but have mostly relied on wrist-worn inertial measurement unit sensors and proximity sensors, which are inadequate for identifying specific locations. This limits their utility for accurate and precise monitoring of behaviors or providing feedback to guide behaviors. A potential clinical application is monitoring body-focused repetitive behaviors (BFRBs), recurrent, injurious behaviors directed toward the body, such as nail biting and hair pulling, that are often misdiagnosed and undertreated. Here, we demonstrate that including thermal sensors achieves higher accuracy in position tracking when compared against inertial measurement unit and proximity sensor data alone. Our Tingle device distinguished between behaviors from six locations on the head across 39 adult participants, with high AUROC values (best was back of the head: median (1.0), median absolute deviation (0.0); worst was on the cheek: median (0.93), median absolute deviation (0.09)). This study presents preliminary evidence of the advantage of including thermal sensors for position tracking and the Tingle wearable device’s potential use in a wide variety of settings, including BFRB diagnosis and management.

## Introduction

Accurate monitoring of hand position with respect to the head has many potential applications, ranging from extended reality and computer gaming to monitoring certain clinical conditions. Body-focused repetitive behaviors (BFRBs) represent a class of potentially useful clinical applications. BFRBs are associated with a broad range of mental and neurological illnesses (e.g., excoriation disorder, trichotillomania, autism, Tourette Syndrome, Parkinson’s Disease)^1,2^, where individuals unintentionally cause physical self-harm through repeated behaviors directed toward the body. Common BFRBs include hair pulling, skin picking, and nail biting, and are often misdiagnosed and undertreated^3^. These symptoms affect at least 5% of the population, and as many as 70% of those with one BRFB will have another co-occurring BRFB^4^. Developmental factors such as age and intellectual disabilities may limit a patient’s awareness of his/her behavior^4^. BFRBs can cause significant distress, impairment and physical health consequences (e.g., pain, disfigurement, infection) and are associated with a sense of diminished control over the behavior^5,6^. As such, it is imperative to establish a reliable means to automatically and objectively identify and monitor BFRBs, especially outside of the clinic.

To approximate ecological BFRB monitoring, conventional methods of position tracking rely on proximity- and inertial measurement unit (IMU) sensor-based measures to identify the position of part of a person’s body (such as a hand) relative to another part of the person (such as the head). The Keen device, a wearable-based tracking method created by HabitAware^7^, is one such attempt to monitor BFRBs. The Pavlok^8^, a wrist-worn device which modifies behavior based on user-induced shocks and feedback, has also been used for BFRB treatment, though it is not specifically designed to do so. Neither of these devices have been the subject of a published peer-reviewed study, so it is unclear how well-suited they are for BFRB monitoring or treatment. The reliance of the Pavlok and Keen devices on IMU sensors may cause difficulty in determining hand position relative to the head because of the lack of head position reference data. A two-device approach, most notably the combination of a bracelet and magnetic necklace^3^, has been used to provide an external reference for head position, producing superior results. However, sensitivity to body movement and user discomfort has made a two-device approach impractical.

The Tingle is a wrist-worn position tracking device designed by the MATTER Lab that passively collects thermal, proximity, and IMU sensor data. The goal of the present study was to assess the efficacy of the Tingle in its ability to distinguish between locations of simulated behaviors, and whether the thermal sensors in the Tingle yield potentially valuable information that may improve BFRB detection and monitoring over proximity and IMU sensor data alone. A long short-term memory (LSTM) neural network was trained using these data to detect when the user’s hand is near one of six target locations on the head.

## Results

### Discriminability

We assessed the degree to which data collected at different locations on the head can be distinguished from one another by calculating a “discriminability distance” measure between each unique pair of target locations on the head (mouth, nose, cheek, eyebrow, top-head, back-head). Data from the proximity and IMU sensors were used to calculate this distance (see Methods). We repeated this analysis with data from all three sensor types, and *the addition of thermal sensor data significantly increased the median discriminability distance between respective targets*, for every target, as shown in Table 1 (paired t-test; p-value ≪ 0.05 for all tests with Bonferroni correction). Across 39 participants, the discriminability distance using all three sensors was greatest between the nose and top of the head (median: 3.32, median absolute deviation: 0.66) and smallest between the nose and cheek (median: 1.25, median absolute deviation: 0.84). We also estimated the null distribution of discriminability distances using permutation testing to shuffle the target labels for each unique pair of targets. Wilcoxon signed-rank testing revealed that the median distance value is significantly greater across all pairs of targets when using data from all three sensor types than when using proximity and IMU sensors (Supplementary Tables 1 and 2).

**Table 1.** Discriminability distance measures, p-values and effect sizes from paired t-tests for the Tingle *with no* thermal sensor data and *with* thermal sensor data (median ± MAD)

| Target pair | Discriminability distance:<br>no thermal | Discriminability distance:<br>yes thermal | p-value | Effect size |
| --- | --- | --- | --- | --- |
| Mouth - Nose | 2.11 ± 1.08 | 3.06 ± 1.04 | 1.29e-13 | 0.90 |
| Mouth - Cheek | 2.30 ± 0.44 | 3.07 ± 0.94 | 4.91e-12 | 1.07 |
| Mouth - Eyebrow | 1.67 ± 1.08 | 2.61 ± 0.75 | 3.05e-11 | 0.97 |
| Mouth - Top-head | 2.32 ± 0.66 | 2.94 ± 0.48 | 3.03e-12 | 0.76 |
| Mouth - Back-head | 2.59 ± 0.57 | 3.31 ± 0.47 | 4.35e-12 | 1.26 |
| Nose - Cheek | 0.49 ± 0.46 | 1.25 ± 0.84 | 3.06e-09 | 0.82 |
| Nose - Eyebrow | 0.85 ± 0.90 | 2.17 ± 0.84 | 1.00e-11 | 1.22 |
| Nose - Top-head | 2.19 ± 0.91 | 3.27 ± 0.91 | 5.52e-13 | 1.06 |
| Nose - Back-head | 2.12 ± 1.12 | 3.32 ± 0.66 | 9.07e-09 | 1.18 |
| Cheek - Eyebrow | 0.77 ± 0.81 | 2.23 ± 1.32 | 5.50e-12 | 1.13 |
| Cheek - Top-head | 2.18 ± 0.67 | 3.22 ± 0.96 | 6.01e-15 | 1.00 |
| Cheek - Back-head | 1.63 ± 1.12 | 3.01 ± 0.72 | 2.83e-11 | 1.09 |
| Eyebrow - Top-head | 1.91 ± 1.01 | 2.71 ± 0.91 | 4.09e-09 | 0.65 |
| Eyebrow - Back-head | 2.33 ± 0.78 | 3.01 ± 0.77 | 1.30e-11 | 0.78 |
| Top-head - Back-head | 2.11 ± 0.90 | 2.88 ± 0.50 | 1.13e-13 | 1.00 |

### Neural Network

[The same data as above were used in this analysis.] To assess the ability of the Tingle to differentiate a hand’s position between target locations on the head, an LSTM neural network^9^ was trained to identify data as on-target (on the correct one of the six targets on the head) or off-target (on one of the remaining five targets, or off of the body). The LSTM network performed this binary classification task for each of the six target regions on the head. This analysis was conducted twice, once without and once with data from the Tingle’s thermal sensors. Accuracy was evaluated using the area under receiver operating curve (AUROC) values, an aggregate measure of specificity and sensitivity^10^, as well as confusion matrices for each binary classification task. Our results shown in Figure 1D demonstrate that the addition of thermal sensor data improves the ability to distinguish between six positions on the head, and does so with median AUROC values > 0.90 for all target locations (Supplementary Table 3). Confusion matrices were assessed for class imbalances in prediction accuracy, but none were identified (Figure 1E).

**Figure 1.**
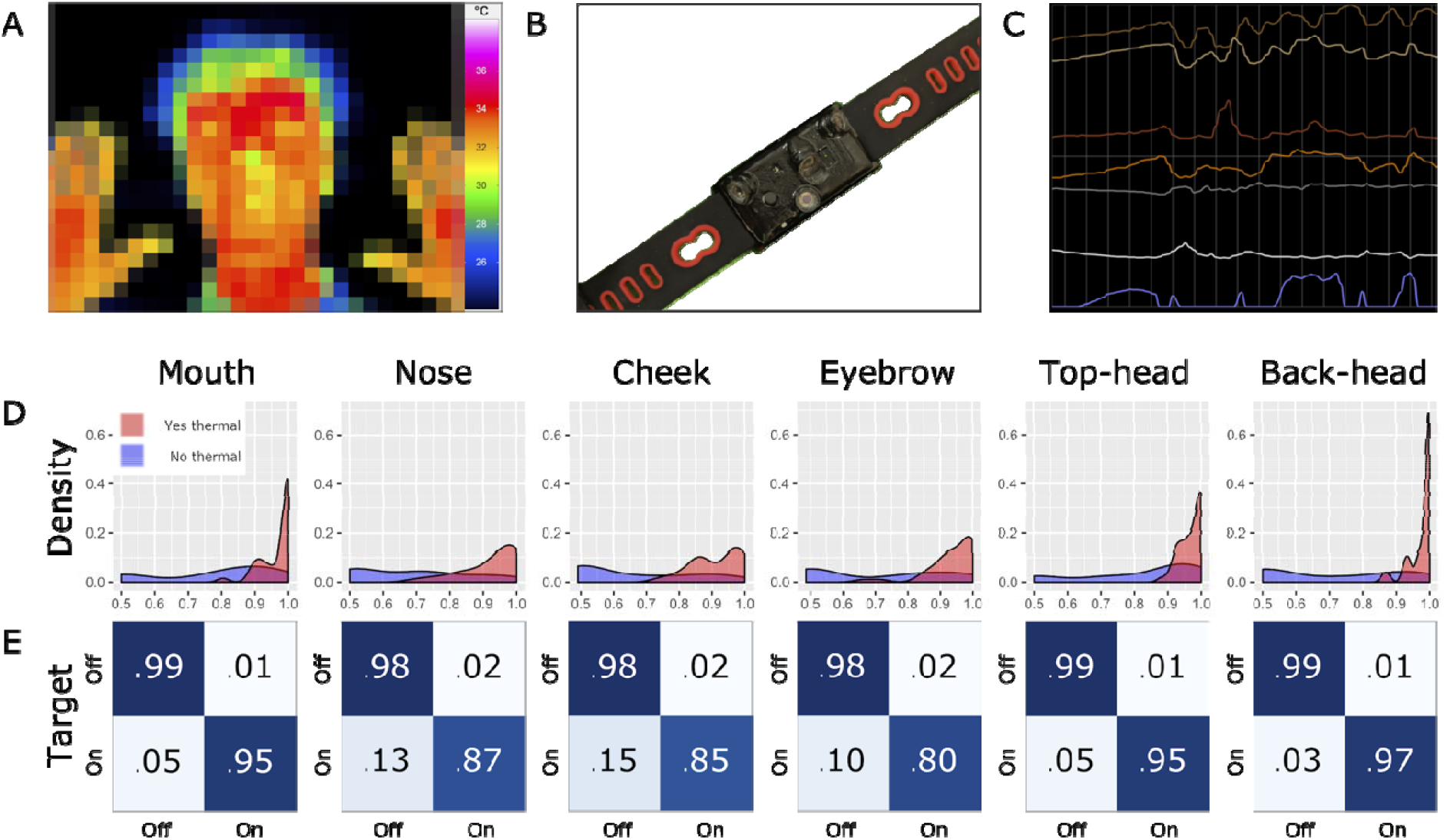
A. Thermal map of the head showing temperature differences (the image was pixelated for this preprint). B. A prototype of the Tingle device, which uses an array of four 1-pixel sensors with different fields of view, a proximity sensor, and two IMU sensors. C. Sample datastream in the Tingle application interface as the user approaches the mouth and hovers around various parts of the head. The top four signals are temperature readings from the four thermopiles, followed by the two IMU sensors, then the proximity sensor in blue. D. LSTM network-based AUROC value distribution for the Tingle per target location on the head. E. Confusion matrices for each location on the head with median values of classifier accuracy across participants.

A general (participant-independent) classifier with the same neural network architecture was also trained using the aggregate data from all participants but one, and then tested on the remaining participant. This leave-one-out approach was applied to create and test 39 general classifiers. All generalized classification accuracy measures were below the median value across individuals but with median AUROC values > 0.80 for all but one (cheek [.75]) target location, as shown in Supplementary Table 3.

## Discussion

In this study, we tested whether thermal sensor data improved the Tingle device’s ability to distinguish between a hand’s position at different locations on the head, for use in detection of a wide range of behaviors, including clinically relevant BFRBs. This investigation was a prerequisite to future studies relevant to distinguishing clinically relevant gestures (BFBRs) from activities of daily living (non-BFRBs). Adding thermal data collected from the Tingle wrist-worn device significantly increased the discriminability distance between targets, and significantly increased the accuracy of a binary classifier based on an LSTM neural network. With thermal data, LSTM neural network model performance has a high degree of accuracy and is comparable across all targets. Without thermal data, the results are more variable, and some classifiers failed to predict any data samples as on-target, resulting in the worst performance AUROC values of 0.5. The general classifier showed promising accuracy measures, demonstrating its potential as a pre-trained model for BFRB detection without individual training.

This study demonstrates that thermal data detected by the Tingle wrist-worn device can help to accurately distinguish between hand locations with respect to the wearer’s head in a controlled setting. This has dramatic consequences for use in different types of hand movement training, in navigation of virtual environments, and in monitoring and mitigating repetitive, compulsive behaviors. In an effort to help guide behaviors, the Tingle can provide haptic feedback (a “tingle”) during detection of a target location. We envision use of thermal sensors in devices like the Tingle helping to train, navigate, and interact in a wide variety of settings.

## Methods

### Data Collection

39 healthy adult employees of the Child Mind Institute or Child Mind Medical Practice were recruited on a volunteer basis. Participants provided written consent with approval from Chesapeake IRB.

Participants were asked to simulate a series of repetitive behaviors, rotating their elbow in a circular motion while their hand was in a fixed position on one of six target locations on the head. Each of the six behaviors was performed for approximately fifteen seconds at a single location on the same side of the head as the dominant hand (the hand wearing the device). Data were collected during each behavior with a sampling rate for this study ranging from 5-7 Hz, which is dynamically adjusted to minimize power consumption. Rotating the elbow provided different orientation information, simulating different approaches to each target location. A web interface designed for data collection (CW) was used by two researchers (JJS and JCC) throughout the experiment. The researchers independently pressed a button on the interface to indicate that the participant’s hand was near the correct location on the head. Data were marked as on-target when both researchers pressed the button during a simulated behavior. Data from off the body were also collected to incorporate information about environmental conditions into the LSTM network.

### Sensors

The Tingle (designed and fabricated by CW) includes a (Kionix KX126) accelerometer^11^, a (STMicroelectronics VL6180X Time-of-Flight Ranging Sensor) proximity sensor^12^, and four (Melexis MLX90615) thermopiles^13^.

### Data Analysis

We de-identified participant data by labeling each participant with a unique index, and z-scaled all data prior to analysis. To determine the discriminability between pairs of target locations on the head, we calculated the median values from the proximity and IMU sensors for a given target location, and computed the Euclidean distance between the pair of vectors of median values corresponding to each pair of target locations. For instance, we isolated data collected near the nose and near the cheek. A vector representing the nose contained the median of the proximity sensor values and median of the IMU sensor values, and a vector representing the cheek contained corresponding median values from the same two sensors in the new location. The resulting pair of vectors was used to calculate the Euclidean distance; this calculation was repeated for each unique pair of target locations on the head. We then constructed a sampling distribution of the distance measure for each of the target pairs by randomly shuffling the target labels (e.g., nose and cheek) 1,000 times and calculating the Euclidean distance between vectors derived from the proximity and IMU sensors. The median Euclidean distance derived from the permutation test was used to provide a baseline measure of the discriminability distance. We conducted a paired t-test test across each of the target pairs using the median values from the sampling distribution and the original distance values (without shuffled labels). We repeated both analyses after including thermal data, by extending the vectors to include median values from the four thermal sensors. To directly compare the discriminability distance between measurements without and with thermal sensor data, we conducted a Wilcoxon signed-rank test across each of the target pairs. Effect sizes were determined by calculating the median difference between data without and with thermal information, then dividing by the median absolute deviation. A three-layer LSTM neural network was trained in Python using the Keras neural network library. The inputs of the LSTM network consist of the z-centered data from the sensors and do not include any additional features. The first two layers consisted of 50 nodes each and the second included a dropout rate of 0.20 to reduce the risk of overfitting. The third layer consisted of a single node using a sigmoid activation function for binary classification. We created training and testing sets with a test size of 25% of the data available. We computed AUROC values and confusion matrices as measures of accuracy at the participant and group level for each of the six target locations on the head.

---

**Supplementary Table 1.**
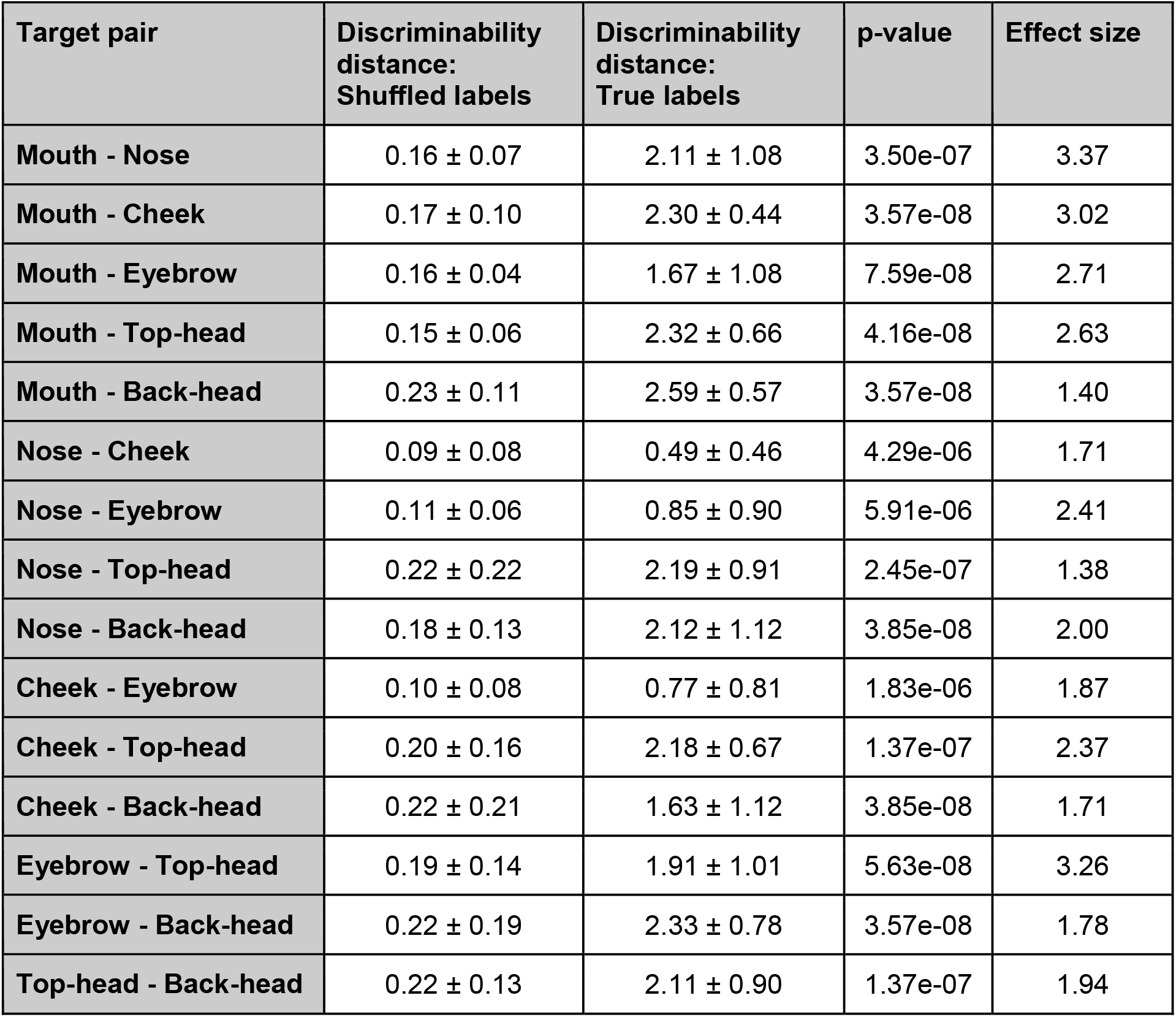
Discriminability distance distribution and p-values from Wilcoxon signed-rank tests for the Tingle *with no* thermal sensor data between the sampling distribution and original distance (median ± MAD)

**Supplementary Table 2.** Discriminability distance distribution and p-values from Wilcoxon signed-rank tests for the Tingle *with* thermal sensor data between the sampling distribution and original distance (median ± MAD)

| Target pair | Discriminability distance:<br>Shuffled labels | Discriminability distance:<br>True labels | p-value | Effect size |
| --- | --- | --- | --- | --- |
| Mouth - Nose | 0.62 ± 0.21 | 3.06 ± 1.04 | 6.07e-08 | 1.91 |
| Mouth - Cheek | 0.71 ± 0.20 | 3.07 ± 0.94 | 4.84e-08 | 1.62 |
| Mouth - Eyebrow | 0.62 ± 0.20 | 2.61 ± 0.75 | 8.18e-08 | 2.04 |
| **Mouth - Top-head** | 0.65 ± 0.11 | 2.94 ± 0.48 | 3.57e-08 | 1.51 |
| **Mouth - Back-head** | 0.76 ± 0.23 | 3.31 ± 0.47 | 3.57e-08 | 1.35 |
| **Nose - Cheek** | 0.52 ± 0.18 | 1.25 ± 0.84 | 3.76e-07 | 1.62 |
| **Nose - Eyebrow** | 0.62 ± 0.38 | 2.17 ± 0.84 | 7.56e-07 | 1.43 |
| **Nose - Top-head** | 1.49 ± 1.14 | 3.27 ± 0.91 | 9.29e-07 | 1.26 |
| **Nose - Back-head** | 0.82 ± 0.41 | 3.32 ± 0.66 | 8.68e-07 | 1.28 |
| **Cheek - Eyebrow** | 0.63 ± 0.32 | 2.23 ± 1.32 | 2.09e-06 | 1.75 |
| **Cheek - Top-head** | 1.05 ± 0.65 | 3.22 ± 0.96 | 9.18e-06 | 1.22 |
| **Cheek - Back-head** | 0.82 ± 0.40 | 3.01 ± 0.72 | 6.54e-08 | 1.31 |
| **Eyebrow - Top-head** | 0.77 ± 0.38 | 2.71 ± 0.91 | 3.31e-06 | 1.32 |
| **Eyebrow - Back-head** | 0.84 ± 0.33 | 3.01 ± 0.77 | 3.57e-08 | 1.51 |
| **Top-head - Back-head** | 0.89 ± 0.40 | 2.88 ± 0.50 | 5.22e-08 | 1.29 |

**Supplementary Table 3.** AUROC value distribution (for individual and general classifiers) and p-values from Wilcoxon signed-rank tests for the Tingle *with no* thermal sensor data and *with* thermal sensor data (median ± MAD)

|  | **Mouth** | **Nose** | **Cheek** | **Eyebrow** | **Top-head** | **Back-head** |
| --- | --- | --- | --- | --- | --- | --- |
| **No thermal** | 0.86 ± 0.13 | 0.69 ± 0.26 | 0.60 ± 0.15 | 0.74 ± 0.36 | 0.91 ± 0.12 | 0.73 ± 0.33 |
| **Yes thermal** | 0.99 ± 0.01 | 0.96 ± 0.07 | 0.93 ± 0.09 | 0.96 ± 0.06 | 0.98 ± 0.06 | 1.00 ± 0.00 |
| **p-value** | 3.20e-07 | 1.14e-07 | 6.15e-08 | 2.48e-07 | 3.65e-07 | 7.25e-08 |
| **Effect size** | 1.27 | 1.37 | 1.93 | 1.66 | 0.94 | 3.49 |
| **General** | 0.89 ± 0.13 | 0.83 ± 0.13 | 0.75 ± 0.15 | 0.84 ± 0.07 | 0.89 ± 0.10 | 0.94 ± 0.08 |

## Contributions

Son, JJ - primary author, data collection and analysis

Clucas, JC - experimental and analytical design, data collection and management, contributed to writing the manuscript

White, C - wearable prototype design and fabrication

Vogelstein, JT - statistical consultation

Milham, M - consultation, reviewed manuscript

Klein, A - experimental and analytical design, contributed to writing the manuscript

## Code Availability

The web applications used to collect data are online at matter.childmind.org/tingle/tingle-min and matter.childmind.org/tingle/tingle-min2, and the code for these sites and analyses are available at github.com/ChildMindInstitute/tingle-pilot-study.

## Data Availability

The datasets analyzed during the current study will be made publicly available at matter.childmind.org/tingle.

## Ethics Statement

This study was carried out in accordance with the recommendations of the Chesapeake IRB with written informed consent from all subjects. All subjects gave written informed consent in accordance with the Declaration of Helsinki. The protocol was approved by the Chesapeake IRB.

## Acknowledgements

The work presented here was primarily supported by gifts to the Child Mind Institute from Phyllis Green, Randolph Cowen, and Joseph Healey. We would like to thank all of our colleagues who participated in this study, and our other colleagues who supported us! JTV would like to acknowledge the Defense Advanced Research Projects Agency (DARPA) Lifelong Learning Machines program. A.Kr. received support from the IDEFI IIFR grant (ANR-2012-IDEFI-04). On behalf of the MATTER Lab, AK would like to thank the Child Mind Institute for supporting the development of the Tingle device.

